# Species detection within the *Echinococcus granulosus sensu lato* complex by novel probe-based Real-Time PCRs

**DOI:** 10.1101/2020.07.24.220756

**Authors:** Pavlo Maksimov, Hannes Bergmann, Marion Wassermann, Thomas Romig, Bruno Gottstein, Adriano Casulli, Franz J. Conraths

## Abstract

Infections with eggs of *Echinococcus granulosus sensu lato* (*s*.*l*.) can cause cystic echinococcosis in intermediate host animals and humans. Upon ingestion of viable eggs, oncospheres hatch from the eggs and subsequently develop into fluid-filled larval cysts, most frequently in the liver or the lungs. The slowly growing cysts progressively interfere with organ function. The risk of infection is determined by the host range of the parasite, its pathogenicity and other epidemiologically relevant parameters, which differ significantly among the five species within the *E. granulosus s*.*l*. complex. It is therefore essential to diagnose the correct species within *E. granulosus s*.*l*. to help understand specific disease epidemiology and to facilitate effective implementation of control measures. For this purpose, simple, fast and cost-effective typing techniques are needed.

We developed quantitative Real-Time Polymerase Chain Reactions (qPCRs) and corresponding sequence-specific hydrolysis DNA probes to target polymorphic regions in the mitochondrial genome of *E. granulosus s*.*l*.. In a single-step typing approach, we distinguished *E. granulosus s*.*l*. members in four epidemiologically relevant subgroups. These were *E. granulosus sensu stricto* (G1, G3), *E. equinus* (G4), *E. ortleppi* (G5) and the *E. canadensis* cluster (G6 to G8 and G10). The technique also allowed identification and differentiation of these species from other *Echinococcus* or *Taenia* taxa for samples isolated from cysts or faeces.

Single-step genotyping techniques for the molecular diagnosis of *Echinococcus* spp. by qPCRs may not only improve diagnostic performance, but also our knowledge on the epidemiology of the parasites and help controlling the various agents of cystic echinococcosis.

## Introduction

Infection with *Echinococcus granulosus sensu lato* (*E. granulosus s*.*l*.) (1) can cause cystic echinococcosis [CE] in animals and humans (2-5). The adult stages of *E. granulosus s*.*l*. are small, segmented tapeworms (Cestoda) that live in the small intestine of their definitive hosts, mostly wild or domestic canids or felids (6, 7). Adult worms release eggs that form the infectious stage for intermediate hosts. These eggs are shed in the faeces of the definitive hosts and are then ingested by intermediate hosts i.e. typically herbivores such as sheep, goats, cattle, deer or camels that are preyed upon by the definitive host (7). After ingestion of viable eggs, oncospheres hatch from the eggs and migrate from the gut into the target organs, mostly liver or lungs (8). Here, the oncospheres convert into the metacestode stage, a slowly growing, fluid-filled cyst containing protoscoleces. This process may progressively interfere with the function of the organs in which the cysts have localised and may lead to clinical or sub-clincal CE in the infected intermediate host (9). Finally, ingestion of fertile cystic material by a definite host, followed by development of the protoscoleces into egg-producing adult tapeworms in their small intestine completes the lifecycle of *E. granulosus s*.*l*. (8).

Previously, *E. granulosus* was considered to be a single species, subdivided into ‘strains’ and genotypes (G1 to G8, G10 and the ‘lion strain’) (4), most of them associated with specific definitive-intermediate host relationships and geographic distribution patterns (7). Currently, some of these taxa are considered distinct species within the *E. granulosus s*.*l*. complex, while others are retained as genotypes within some of these species. Thus, *E. granulosus sensu stricto* (*s*.*s*.) is composed of genotypes G1 and G3, while G2 is no longer considered a distinct genotype (10). In addition, the G1-G3 cluster includes other closely related haplotypes, which – depending on the definition – may or may not belong to these genotypes. These are named Gx in this study. *E. granulosus s*.*s*. is mainly associated with sheep as its intermediate host and causes the largest number of human cases worldwide. Second, *E. equinus* (formerly genotype G4) is associated with horses and rarely, if ever, causes human disease. Third, *E. ortleppi* (formerly genotype G5), associated with cattle, seems to be of low pathogenicity to humans, while, fourth, *E. canadensis* (composed of genotypes G6, G7, G8 and G10) is mainly associated with cervids, camels, goats and pigs and causes the second largest number of human patients; it may have to be subdivided into two or more species and is best referred to as ‘*E. canadensis*-cluster’. Lastly, *E. felidis* (formerly ‘lion strain’) is a wildlife parasite from sub-Saharan Africa, which seems to be present only where lions still exist; no human cases have been reported so far. Apart from these five species, a yet unnamed taxon (G-Omo), related to, but not belonging to the G1-3 cluster of *E. granulosus s*.*s*., has been described from a human patient in southern Ethiopia (4, 11-15).

Accurate identification and reporting to the species level within *E. granulosus s*.*l*. is critical to prevent or control parasite spread and further accidental ingestion by humans. With possible exception of *E. felidis*, all species / genotypes are zoonotic and can cause serious CE in humans, who act as aberrant, dead-end intermediate hosts. Identifying the individual species within *E. granulosus s*.*l*. is also necessary for detailed epidemiological understanding of the disease in its geographic distribution, for surveillance purposes, and ultimately for implementing effective prevention and control measures. This is particularly important in endemic areas, where several *Echinococcus* species and other taeniid cestodes coexist. This differentiation is also important to assess the different zoonotic potential of the various species (16) in intermediate and aberrant hosts at risk (16, 17). According to a recent expert consensus, specific differentiation of the *E. granulosus s*.*l*. complex should be done with any kind of sample whenever technically feasible (1).

Methods used for the diagnosis of *Echinococcus* infections in definitive and intermediate hosts include classical parasitological techniques such as microscopic examination of intestinal scrapings, counting procedures and sedimentation and flotation of faecal samples (18-22) as well as molecular techniques such as polymerase chain reaction [PCR] which are particularly useful for species or genotype differentiation (16, 17). The traditional parasitological methods are important for the preparation and pre-selection of sample material, but generally do not provide sufficient acuity for species differentiation. Several PCR protocols for detection and differentiation of species within *E. granulosus s*.*l*. have been published (23-26). However, most of these protocols are based on conventional PCR techniques and either require an additional sequencing step for unambiguous determination, or are limited to certain *Echinococcus* species, taxa or genotypes. Therefore, a tool for simultaneous testing for and typing of species within *E. granulosus s*.*l*. in a parallel manner would be a helpful addition to the diagnostic toolbox. Such a tool would enable reliable, simple, fast and affordable diagnosis to the species level (*E. granulosus s*.*s*., *E. equinus, E. ortleppi* [G5]) and *E. canadensis*).

Here we show that sequence-specific DNA probe-based quantitative PCR [qPCR] amplification of polymorphic target regions can be used as a single-step diagnostic tool for the differentiation of the four most important *Echinococcus* species that cause cystic echinococcosis. Diagnostic DNA probe qPCR tests were successfully applied, also together with an internal control in duplex or triplex format to identify the single species derived from cystic material or faecal matter. We anticipate that this tool will not only make diagnosis easier and faster, but would also be of use for epidemiological studies, effective prevention planning and control programs (16, 17).

## Materials and Methods

### Primer and DNA probe design for qPCRs

For a bioinformatic prediction of suitable regions in the mitochondrial genome of *Echinococcus* spp., relevant sequences were downloaded from the public NCBI database at https://www.ncbi.nlm.nih.gov/nucleotide/. Accession numbers are listed in Table 1 and sequence analysis is summarised in Figure 1. Mitochondrial genomes were aligned using the “MAFFT” algorithm embedded in the Geneious Prime^®^ (Version 2019.2.3) software suite. The program was used with default settings, whereby selection of the appropriate algorithm for the alignment was set to automatic. The Primer 3 algorithm, embedded in Geneious Prime^®^, was then applied to all aligned sequences to predict potential targets for qPCR-based diagnosis. Thereafter, the *in silico* predicted qPCR targets were further analysed to find suitable polymorphic regions that might allow differentiating *E. granulosus s*.*l*. species of interest from others within the *E. granulosus s*.*l*. complex. To validate selected primer sequences *in vitro*, their performance was assessed in a SYBR® Green qPCR assay as described below.

**Table 1:**
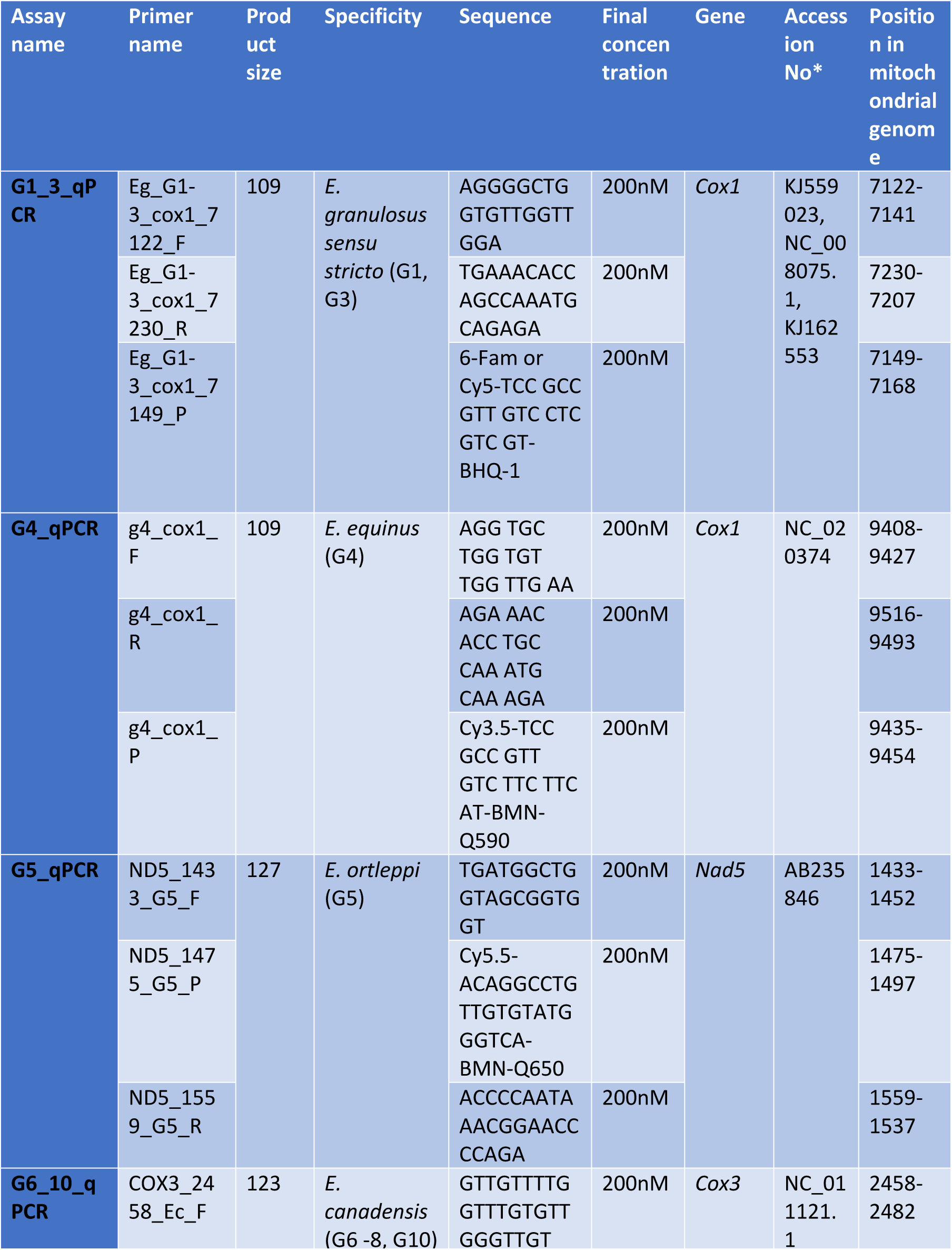
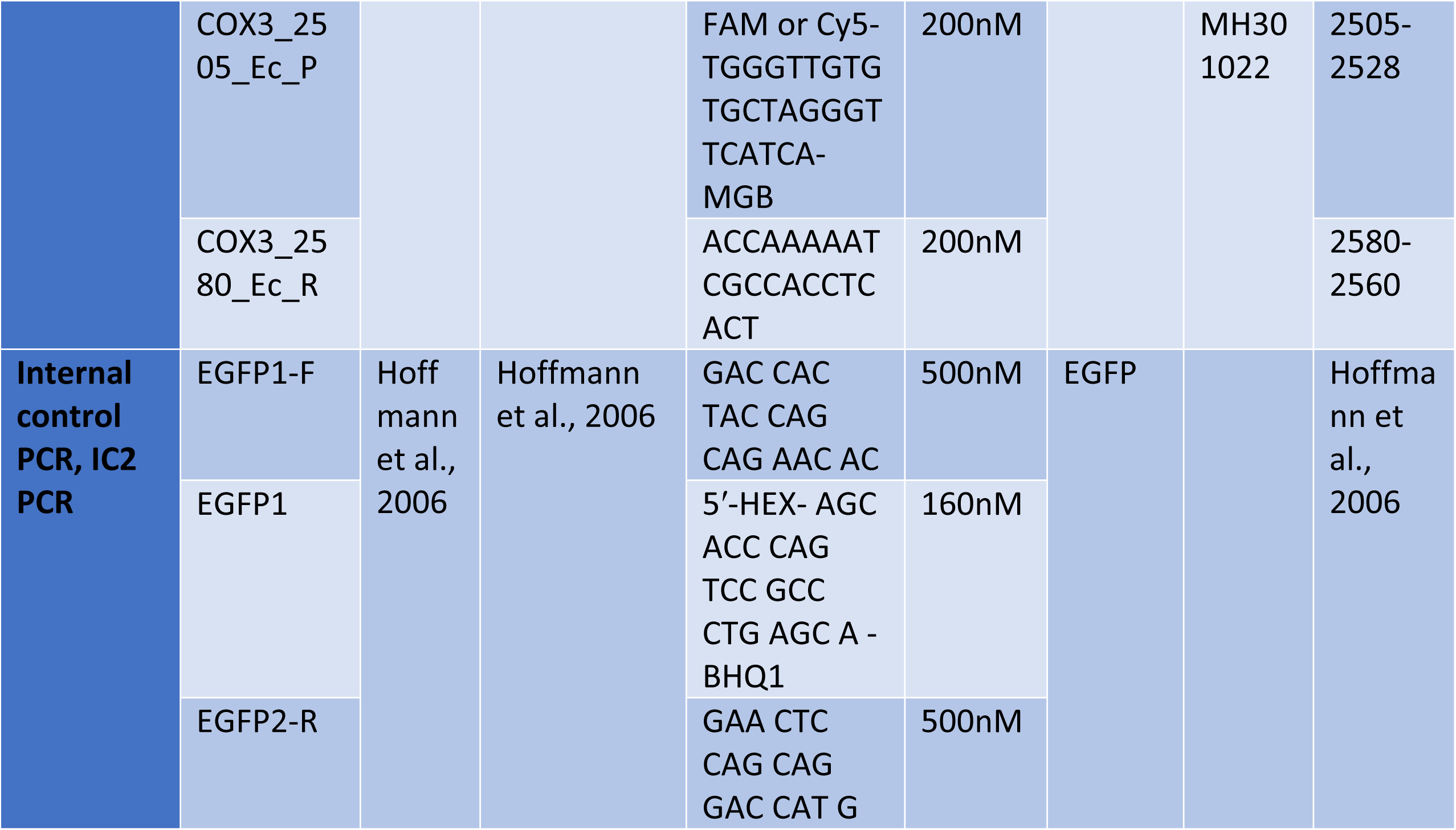
Identified primer and probe sequences for detection and typing of *Echinococcus granulosus sensu lato* species by TaqMan® qPCR.

**Figure 1:**
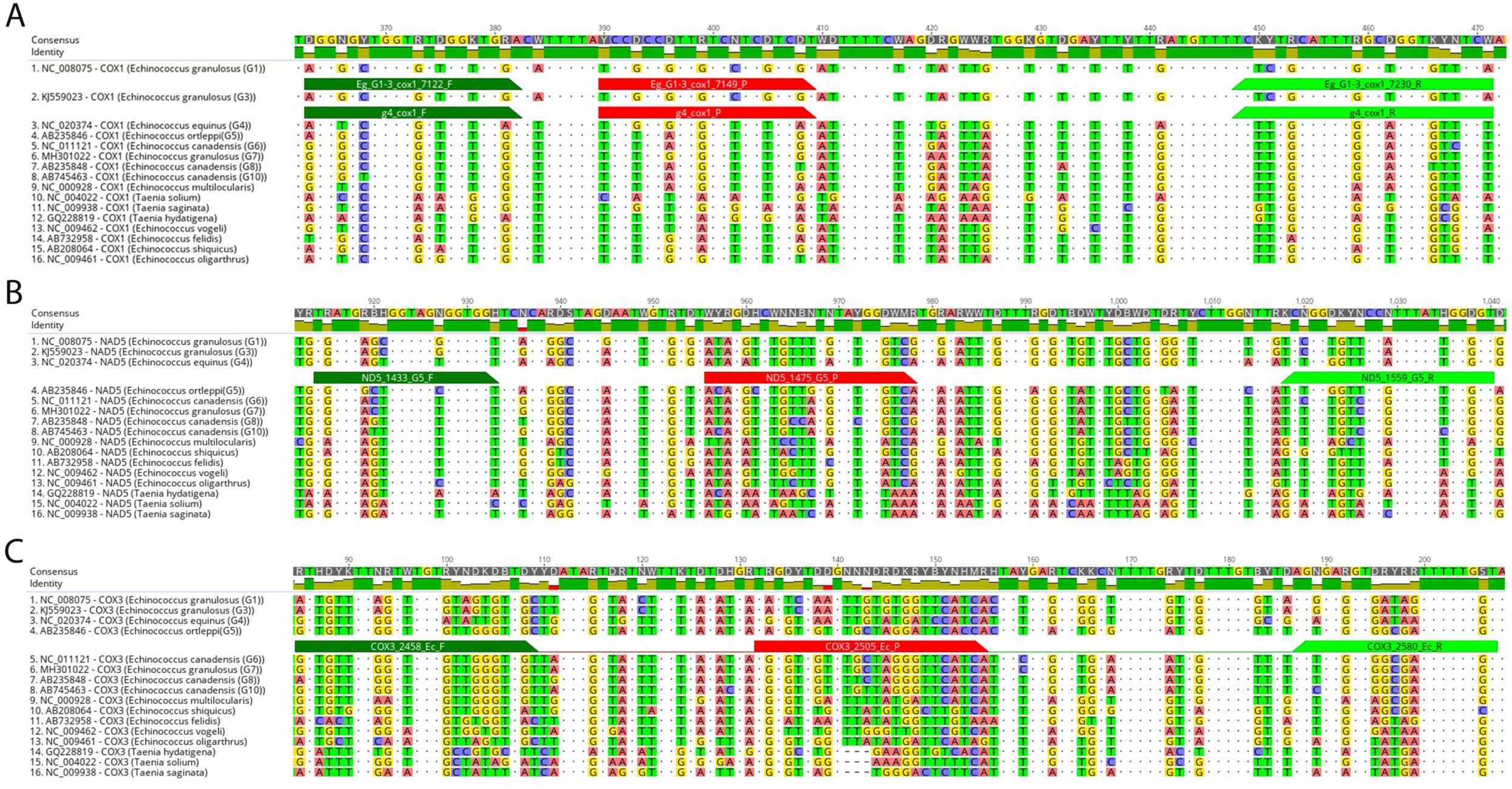
Location of *Echinococcus granulosus* sensu lato polymorphic genome regions within COX1, COX3 and NAD5 genes used for primer and probes design. *In-silico* identification of polymorphic regions in aligned mitochondrial DNA sequences to differentiate the *E. granulosus sensu lato* species *E. granulosus sensu stricto* (G1-3): **A**, *E. equinus* (G4): **A**, *E. ortleppi* (G5): **B** and *E. canadensis* (G6-8, G10): **C** from other *Echinococcus* spp. and *Taenia* spp. Differences between aligned sequences are highlighted by different background colours of respective nucleotides. Dots represent nucleotides identical in all aligned sequences. Dark-green bars show the positions of the respective forward primer, red bars annotate the positions of the respective probes, and light-green bars indicate the positions of the reverse primer. The name of each aligned sequence consists of NCBI nucleotide data base accession number, followed by gene name and the name of *Echinococcus* species. G: genotype

### Reference DNA samples and faecal spiking

To develop and validate the qPCR, we utilised a panel of reference DNA samples from members of the *E. granulosus s*.*l*. complex and related cestode species as outgroup controls. We included *E. granulosus s*.*s*. (G1-3), *E. equinus* (G4), *E. ortleppi* (G5) and genotypes of the *E. canadensis* cluster (G6, G7, G8 and G10) (Table 2). To verify the identity of the taxa in the reference material, we genotyped all DNA samples by conventional PCR (Figure 2) as described in this manuscript and shipped the obtained amplicons to Eurofins Genomics (Ebersberg, Germany) for Sanger sequencing. To minimise the cross-contamination risk among reference samples, the DNA quantity was reduced by diluting samples 1:100. These dilutions were used as working samples in all experiments. To test the analytical sensitivity and efficiency of the TaqMan^®^ qPCR assays, DNA samples were further diluted in ten-fold steps up to 1:10^−7^ in Tris-EDTA buffer with 2 µg/µl Bovine Serum Albumin [BSA] (Carl Roth, Karlsruhe, Germany) (27). As faecal matter is known to contain PCR inhibitory components (28), spiked faecal samples from a confirmed *E. multilocularis*-free red fox (*Vulpes vulpes*) were prepared. To test whether *E. granulosus s*.*l*. species could be differentiated in DNA samples extracted from faecal samples, 200 mg of faeces were spiked with reference DNA that had been serially diluted in ten-fold steps up to 1:10^−4^. Each sample was prepared in duplicate. Spiked faecal samples were then processed using the ZR Faecal DNA MiniPrep^™^ kit (Zymo Research, Freiburg, Germany) to carry out DNA extraction as recommended in the kit manual.

**Table 2:** Reference DNA sample characteristics.

| Species | Genotype | Animal origin | Geographical origin |
| --- | --- | --- | --- |
| *E. granulosus s.s.* | G1 | Cattle | Armenia |
| *E. granulosus s.s.* | G3 (,G2') | Sheep | Armenia |
| *E. granulosus s.s.* | G3 | Sheep | France |
| *E. granulosus s.s.* | Gx | Cattle | Armenia |
| *E. granulosus s.s.* | Gx | Schaf | Ethiopia |
| *E. equinus* | G4 | Zebra | Namibia |
| *E. equinus* | G4 | NA | NA |
| *E. ortleppi* | G5 | NA | NA |
| *E. ortleppi* | G5 | Cattle | Kenya |
| *E. ortleppi* | G5 | Cattle | Zambia |
| *E. canadensis* | G6 | Camel | Kenya |
| *E. canadensis* | G6 | Camel | Kenya |
| *E. canadensis* | G7 | NA | NA |
| *E. canadensis* | G8 | NA | Russia |
| *E. canadensis* | G10 | NA | NA |
| *E. canadensis* | G10 | Deer (Cervus sp.) | Russia |
| *E. vogeli* |  | NA | NA |
| *E. felidis* |  | Warthog | Namibia |
| *E. cf. granulosus* | G-Omo | Human | Ethiopia |
| *T. saginata* |  | Cattle | Ethiopia |
| *T. hydatigena* |  | Dog | Ethiopia |
| *E. multilocularis* |  | Red fox | Germany |

*s.s.*: *sensu stricto*

**Figure 2:**
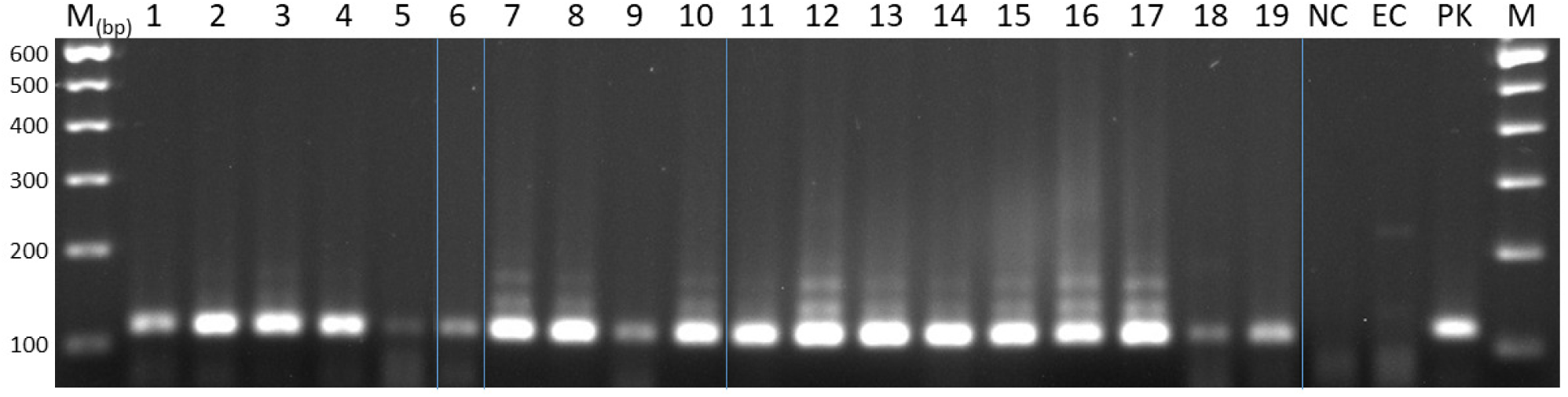
Reference DNA amplification of specific PCR products of *Echinococcus granulosus sensu lato*. PCR products from reference DNAs obtained from *Echinococcus granulosus sensu lato* species amplified by specific PCR (Cest4, Cest5 (Trachsel et al., 2007)). 1: *E. granulosus sensu stricto* (G1), 2: *E. granulosus s*.*s*. (G3 [“G2”]), 3: *E. granulosus s*.*s*. (G3), 4-5: *E. granulosus s*.*s*. (Gx), 6-7: *E. equinus* (G4), 8-10: *E. ortleppi* (G5), 11-12: *E. canadensis* (G6), 13: *E. canadensis* (G7), 14: *E. canadensis* (G8), 15-16: *E. canadensis* (G10), 17: *E. vogeli*, 18: *E. felidis*, 19: *E. cf. granulosus* (G-Omo). M: 100 base pair (bp) marker. NC: non-template control, EC: negative extraction control. PK: positive control. Blue vertical lines: Gel spliced to clean up image and for labeling purposes

### Conventional PCR

To confirm the identity of the respective *Echinococcus* spp. in reference DNA samples, conventional PCR was performed as described (23, 28). In brief, the following primers were used to amplify *E. multilocularis* DNA sequences (H15 F [5-CCATATTACAACAATATTCCTATC-3], EM-H17 R[5-GTGAGTGATTCTTGTTAGGGGAAG-3]), *E. granulosus s*.*l*. DNA (Cest4 F[5-GTTTTTGTGTGTTACATTAATAAGGGTG-3], Cest5 R[5-GCGGTGTGTACMTGAGCTAAAC-3]) and *Taenia* spp. DNA (Cest3 F[5-YGAYTCTTTTTAGGGGAAGGTGTG-3], Cest5 R[5-GCGGTGTGTACMTGAGCTAAAC-3]) (23, 29). PCR was performed in a total volume of 25 µl per sample with 2.5 µl 10x PCR Rxn Buffer (Invitrogen Platinum^®^Taq Polymerase, Invitrogen GmbH, Darmstadt, Germany), 100 pmol/µl of each primer (synthesized at Eurofins Genomics,Ebersberg, Germany), 12.5 mM dNTPs, 50 mM MgCl_2_ (InvitrogenPlatinum^®^Taq Polymerase), 5 U/µl Taq Polymerase (Invitrogen Platinum^®^Taq Polymerase, Qty. 300 Rxn) and 2.5 µl DNA template. The PCR was performed in a Bio-Rad C1000 Thermal Cycler Detection System (Hercules,Bio-Rad Laboratories GmbH, Munich, Germany) applying the following cycle: 94 °C for 3 min, thereafter 40 cycles with 94 °C for 30 s, 55 °C for 30 s and 72 °C for 60 s. PCR products were separated by agarose gel electrophoresis in 2 % gels (Biozym^®^LEAgarose, Hessisch Oldendorf, Germany) with 0.2 % ethidium bromide.

### Generic quantitative PCR (SYBR^®^ Green)

To test the specificity of *in silico* selected primers, a generic quantitative PCR was carried out with a panel of reference DNAs. Primer and probes were obtained from Metabion (Planegg, Germany) (Table 2). The SYBR^®^ Green qPCR was performed in a total volume of 20 µl. For the PCR reaction mixture, 10 µl SsoAdvanced universal SYBR^®^ Green supermix (2x) (Biorad Laboratories GmbH, Munich, Germany), 10 pmol of each primer (forward and reverse), ultrapure nuclease-free water (Sigma-Aldrich, Missouri, USA) and 5 µl of template DNA were used. The cycling conditions in the SYBR^®^ Green real-time PCR were 98.0 °C (3 min, activating of Taq polymerase), followed by 40 cycles at 95.0 °C for 15 s and at 60.0 °C for 30 s. After each cycle, the light emitted by the fluorophore was measured. The melting curve was constructed from 65 °C to 95 °C at 0.5 °C increments with a dwell time of 5 s at each temperature. Real-time PCR results were analysed using the CFX Maestro software suite (Version: 3.1.15; Biorad Laboratories GmbH, Munich, Germany).

### Sequence-specific DNA probe based quantitative PCR (TaqMan^®^) and internal control

TaqMan^®^ qPCR was performed as previously described (28, 30). In brief, the total reaction volume of 25 µl per sample included 12.5 µl TaqMan^®^ qPCR master mix (QuantiTect^®^ Multiplex PCR NoROX Kit (QIAGEN, Hilden, Germany), 1.25 µl of the respective primer/probe mix (Table 2) (200 nM of each primer (forward and reverse) together with 200 nM hydrolysis probe), 6.25 µl ultrapure nuclease-free water (Sigma-Aldrich, Missouri, USA) and 5 µl DNA template. Primers and probes were obtained from Eurofins Genomics (Ebersberg, Germany). The qPCR was carried out in a Bio-Rad CFX 96 Real-Time Detection System (Hercules, Bio-Rad Laboratories GmbH, Munich, Germany) using the following thermal profile: 50 °C for 2 min, an initial denaturation step at 95 °C for 15 min and 45 amplification cycles of 94 °C for 60 s followed by annealing and elongation at 59 °C for 1 min. When the reference DNA-spiked faecal samples were analysed by TaqMan^®^ qPCR, a known quantity of heterologous plasmid DNA containing the enhanced green fluorescent protein [EGFP] gene (31) was included as an internal control [IC] in the qPCR mixture to detect potential inhibition. IC plasmids were added to each TaqMan^®^ qPCR reaction mix along with the primers EGFP1-F, EGFP1-R and the probe EGFP1 (Table 2). The amount of the IC DNA added to each reaction was adjusted to result in a Cq value of about 30 in the respective qPCR tests. Each sample was tested by adding 12.5 μl QuantiTect^®^ Multiplex PCR NoROX Kit (200 × 50 μl reactions, QIAGEN, Hilden, Germany), 1 µl IC-DNA, 1.25 µl of the respective primer/probe mix to detect *Echinococcus* spp. DNA, 0.3 µl EGFP primer/probe mix and 4.95 µl ultrapure nuclease-free water (Sigma-Aldrich, Missouri, USA). Due to a cross-reaction of the G6_10_qPCR primers with *E. ortleppi* (G5) DNA, this qPCR reaction had to be modified into a duplex format by combining it with G5_qPCR primers and probe. This duplex assay was further combined with the EGFP-PCR in a triplex format to test DNA samples extracted from faecal samples. The concentrations of the corresponding primers and probes are shown in the Table 2. Reactions were run in a total volume of 25 μl, which included 5 μl template DNA extracted from faecal samples. The duplex and triplex TaqMan^®^ qPCRs were also carried out in the Bio-Rad CFX 96 Real-Time Detection System (Hercules, Bio-Rad Laboratories GmbH, Munich, Germany), using the same thermal program as described above. Emitted fluorescence was measured at the end of every cycle. A negative extraction control that had been used in parallel throughout the DNA extraction process, a negative PCR control sample (sterile deionized water) and a positive control were included in all qPCR runs.

### Sequencing of qPCR products

To verify qPCR amplicon sequences, purified amplicons from each single-plex TaqMan^®^ qPCR were cloned into a plasmid vector using the pGEM^®^-T Easy Vector System I kit (Cat.# A1360, Promega, Walldorf, Germany) and One Shot^®^ TOP10 (Thermo Fisher Scientific, Waltham, MA, USA)) chemically competent *Escherichia coli* according to the manufacturer’s instructions. Each plasmid vector DNA was then extracted from cultivated *E. coli* with the QIAprep Spin Miniprep Kit (QIAGEN, Hilden, Germany) according to the manufacturer’s instructions and subsequently sequenced using the BigDye Terminator v1.1 Cycle Sequencing Kit and an ABI 3130 capillary sequencer (Thermo Fisher Scientific, Langenselbold, Germany). Sequences were aligned and assembled with the Geneious Prime^®^ (2019.2.3) software package. The concentration of plasmid DNA containing the respective qPCR-specific products was determined by the Nanodrop technology (Thermo Fisher Scientific, Waltham, MA, USA)). The plasmid DNA copy number was estimated using an online tool (Plasmid DNA copy calculator; https://www.thermofisher.com/de/en/home/brands/thermo-scientific/molecular-biology/molecular-biology-learning-center/molecular-biology-resource-library/thermo-scientific-web-tools/dna-copy-number-calculator.html).

## Results

### Primer selection for the amplification of *E. granulosus* s. l. species sequences

To design PCR primers and TaqMan^®^ PCR probes that allow diagnostic differentiation between the targeted species, mitochondrial genome sequences were bioinformatically analysed (Figure 1). The mitochondrial genomes of species and intraspecific genotypes from the *E. granulosus s. l*. complex were screened for polymorphic regions that could be used as targets for specific PCR primer pairs and corresponding probes. Identified regions and verified primers and probes for all four species, *E. granulosus s*.*s*. (G1-G3), *E. equinus* (G4), *E. ortleppi* (G5), and *E. canadensis* (G6-10), are shown in Table 1 and Figure 1.

We used reference samples of known tapeworm species and genotypes to test the newly identified primer pairs and probes. To ensure the identity of the reference material, tapeworm specific DNA was amplified by conventional PCR and the resulting amplicons were then analysed by Sanger sequencing. The conventional PCR and sequencing results confirmed the identity of the DNA reference samples in all cases (Figure 2, sequencing results not shown).

The bioinformatically selected primer pairs were then examined by SYBR^®^ green qPCR to test if they specifically amplified sequences from the genome of the targeted species. We found that the selected primer pairs for *E. granulosus s. s*. (G1-G3) and *E. equinus* (G4) specifically amplified their respective reference DNA and generated amplicons that were distinguishable by their melting peak temperatures from amplicons generated using samples with DNA from closely related *Echinococcus* spp. and *Taenia* spp. (Figure 3A and B). Furthermore, with the *E. granulosus s*.*s*. (G1-G3) primers, the melting temperature of 81 °C was distinguishable from the melting temperatures observed in amplicons generated with the same primer pair in reference samples of *E. equinus* (G4), *E. ortleppi* (G5) and *E. canadensis* (G6-10) at 79-80 °C (Figure 3A). The selected primer pair designed for *E. equinus* (G4) generated a specific PCR product and did not cross-react with samples of the other *E. granulosus s*.*l*. species or *E. multilocularis* (Figure 3B, Table 3).

**Table 3:** Specificity test results of TaqMan® qPCR assays.

| Reference DNA |  | qPCR |  |  |  |
| --- | --- | --- | --- | --- | --- |
| Species | Genotype | G1_3_qPCR | G4_qPCR | G5_10_qPCR | G5_10_qPCR |
| *E. granulosus*<br>s.s. | G1 | 31.65 | No Cq | No Cq | No Cq |
| *E. granulosus*<br>s.s. | G3 (G2') | 28.53 | No Cq | No Cq | No Cq |
| *E. granulosus*<br>s.s. | G3 | 27.68 | No Cq | No Cq | No Cq |
| *E. granulosus*<br>s.s. | Gx | 36.90 | No Cq | No Cq | No Cq |
| *E. granulosus*<br>s.s. | Gx | 37.46 | No Cq | No Cq | No Cq |
| *E. equinus* | G4 | No Cq | 39.06 | No Cq | No Cq |
| *E. equinus* | G4 | No Cq | 24.02 | No Cq | No Cq |
| *E. ortleppi* | G5 | No Cq | No Cq | 27.53 | 28.02 |
| *E. ortleppi* | G5 | No Cq | No Cq | 35.96 | 35.54 |
| *E. ortleppi* | G5 | No Cq | No Cq | 26.02 | 26.54 |
| *E. canadensis* | G6 | No Cq | No Cq | No Cq | 37.56 |
| *E. canadensis* | G6 | No Cq | No Cq | No Cq | 28.37 |
| *E. canadensis* | G7 | No Cq | No Cq | No Cq | 25.57 |
| ***E. canadensis*** | G8 | No Cq | No Cq | No Cq | 29.6 |
| ***E. canadensis*** | G10 | No Cq | No Cq | No Cq | 27.11 |
| ***E. canadensis*** | G10 | No Cq | No Cq | No Cq | 21.0 |
| ***E. vogeli*** |  | No Cq | No Cq | No Cq | No Cq |
| ***E. felidis*** |  | No Cq | No Cq | No Cq | No Cq |
| ***E. cf. granulosus*** | G-Omo | No Cq | No Cq | No Cq | No Cq |
| ***T. saginata*** |  | No Cq | No Cq | No Cq | No Cq |
| ***T. saginata*** |  | No Cq | No Cq | No Cq | No Cq |
| ***T. hydatigena*** |  | No Cq | No Cq | No Cq | No Cq |
| ***E. multilocularis*** |  | No Cq | No Cq | No Cq | No Cq |
| ***T. hydatigena*** |  | No Cq | No Cq | No Cq | No Cq |

s.s.: *sensu stricto*

**Figure 3:**
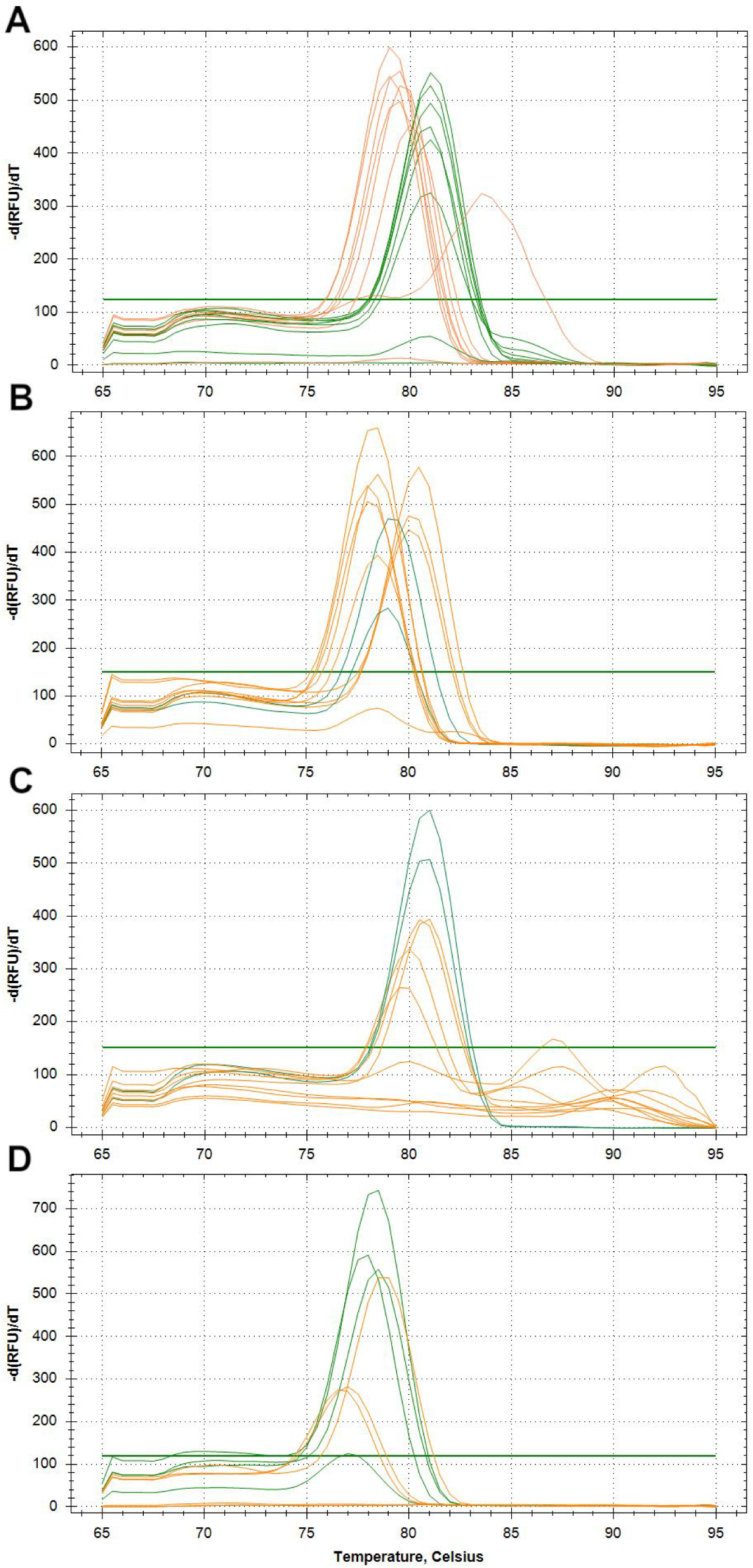
Quantitative SYBR^®^ green PCR melting curves of *Echinococcus* reference DNA samples analysed with newly identified primer pairs. Green lines indicate peaks from targeted reference DNA. Orange lines indicate peaks from not-targeted reference DNAs. A: primer for detection of *Echinococcus granulosus sensu stricto* (G1-G3) DNA. B: primer for detection of *E. equinus* (G4) DNA. C: primer for detection of *E. ortleppi* (G5) DNA. D: Primer for detection of *E. canadensis* (G6-8,G10) DNA

By contrast, primers designed to amplify sequences of *E. ortleppi* (G5), targeting a polymorphic sequence in *Nad5*, also yielded non-specific PCR products in the *E. canadensis* (G6-10) reference samples and vice versa, i.e. primers designed to generate *E. canadensis* (G6-10) amplicons from *Cox3* amplified also *E. ortleppi* (G5) reference DNA. Whilst the selected primer pairs amplified PCR products in *E. granulosus* DNA samples, the resulting melting curves of all of the products were hardly distinguishable at the observed temperatures of 78 °C to 78.5 °C for *E. canadensis* (G6-10) and 79 °C for *E. ortleppi* (G5) (Figure 3C and D). Nevertheless, the primers were selected for further testing in the TaqMan qPCR with their corresponding probes.

Additional primer and probe combinations that were initially selected *in silico*, but did not satisfactorily amplify sequences from the targeted species during SYBR^®^ green, or in further TaqMan^®^ qPCR experiments, are shown in the supplementary Table S1.

In summary, amplicon generating primer pairs were identified for all targeted species (Table 1). Conventional SYBR^®^ green qPCR was sufficient to differentiate *E. granulosus s. s*. (G1-G3) and *E. equinus* (G4) using primers targeting polymorphic regions in *Cox1* (Table 1) based on the generated amplicon alone. Using the same method, primers targeting regions in *Nad5* in *E. ortleppi* (G5) and in *Cox3* in *E. canadensis* (G6-8, G10) amplified successfully, but cross-reacted and required additional testing to separate the respective genotypes.

### Detection of *Echinococcus* species by TaqMan^®^ quantitative PCRs

To increase the specificity of *Echinococcus* spp. target DNA detection, the four selected primer pairs were combined with amplicon-specific TaqMan^®^ DNA probes (Table 1), and used to analyse serially diluted DNA reference samples (Figure 4). TaqMan^®^ qPCR assays for *E. granulosus* s. s. (G1-G3) and *E. equinus* (G4) specifically identified target DNA in the corresponding reference samples. The selected probes for both subgroups did not bind DNA from the other *E. granulosus s*.*l*. species, from *E. multilocularis* spp. or from *Taenia* spp. (Table 3). Primers and probe for detecting *E. granulosus s*.*s*. (G1-G3) also amplified DNA from the G_x,_ strain sample, indicating that the assay is also applicable to other genotypes of *E. granulosus s*.*s*..

**Figure 4:**
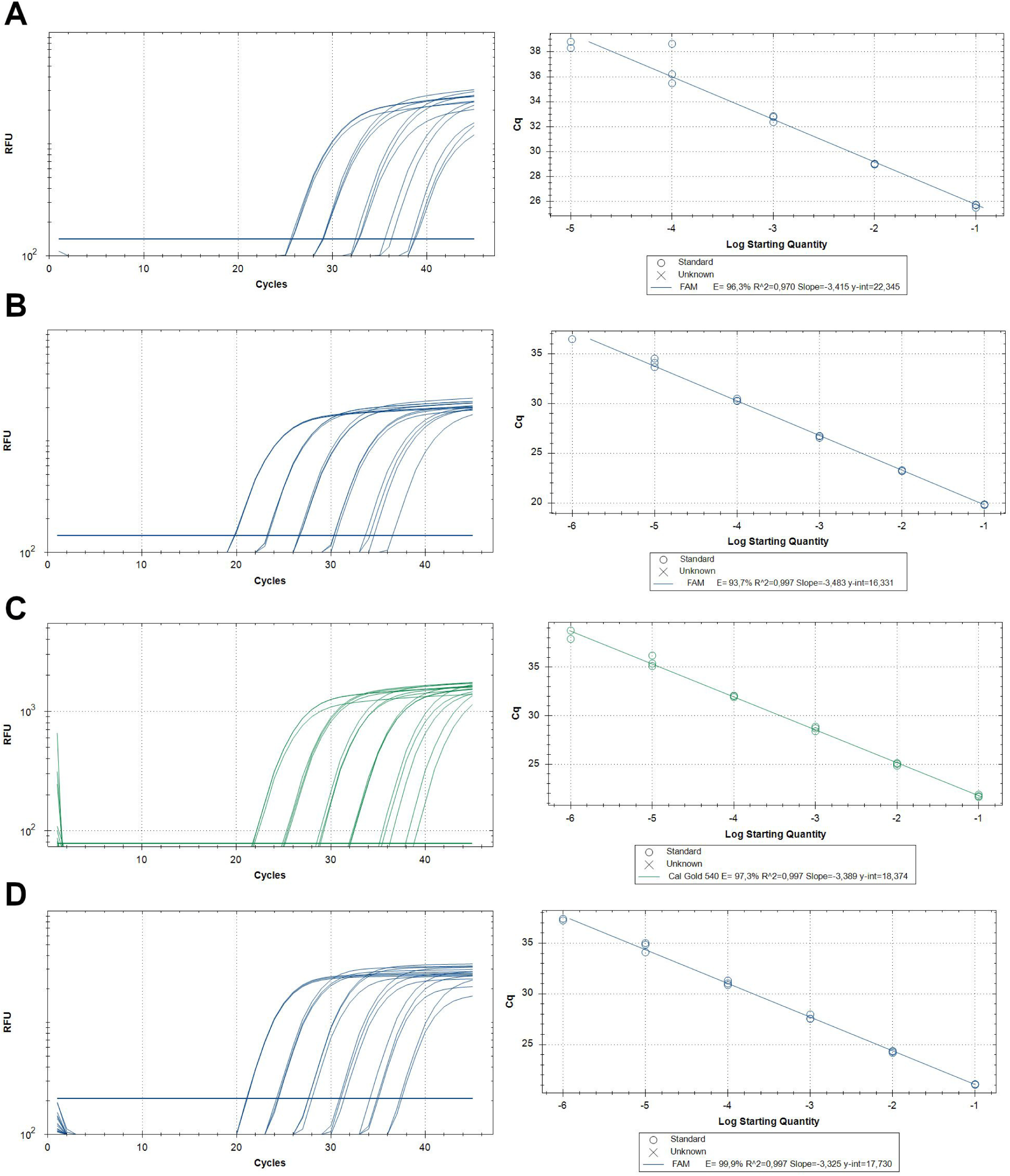
Fluorescence curves for developed TaqMan® qPCRs of serially diluted *Echinococcus* reference DNA samples. **A**: G1_3_qPCR: *E. granulosus s*.*s*., **B**: G4_qPCR: *E. equinus*, **C**: G5_10_qPCR: *E. ortleppi*, and **D**: G5_10_qPCR: *E. canadensis*. RFU: relative fluorescence units. Cq: quantification cycle.

Addition of TaqMan^®^ probes to qPCR reactions with selected G5_qPCR primer pairs for *E. ortleppi* (G5) also showed specific amplification of the DNA target region. No cross-reactions were recorded with DNA samples from the remaining *Echinococcus* spp. and *Taenia* spp. (Table 3). Thus, it appears that the addition of sequence-specific DNA probes ameliorated the cross-reactivity with the *E. canadensis* cluster observed in the qPCR system.

TaqMan^®^ qPCR with the G6_G10_qPCR primer set targeting the *E. canadensis* (G6-8, G10) cluster resulted in amplification of a probe-binding PCR product (Figure 4C and D). However, this assay also cross-reacted with the *E. ortleppi* (G5) DNA sample and thus did not allow differentiation of these two species (Table 3). To distinguish *E canadensis* (G6-8, G10) from G5 samples, we combined the G5_qPCR with the G6_G10_qPCR primer pairs and probes in a TaqMan^®^ qPCR (G5_G10_qPCR) duplex format. With this format only the probe-binding product for *E. canadensis* (G6-8, G10) would be amplified and detected if *E. canadensis* was present in the diagnostic sample. Whereas, if *E. ortleppi* (G5) were present in the diagnostic sample, the probe-binding products for both *E. ortleppi* (G5) and *E. canadensis* (G6-8, G10) genotypes would be detected, as the primer and probe for *E. canadensis* (G6-8, G10) also amplified and detected *E. ortleppi* (G5). Therefore, the duplex TaqMan^®^ qPCR format of G5_G10_qPCR primer and probes allowed diagnostic differentiation of *E. ortleppi* (G5) from the *E. canadensis* (G6-8, G10) cluster (Table 3).

To characterise the diagnostic TaqMan^®^ qPCR assays further, the analytical sensitivity and efficiency of the qPCR reactions for each subgroup were determined by testing DNA extracted from clinical samples as cloned PCR products (plasmid DNA) (Table 4, Figure 4 and Figure 5). The efficiency of TaqMan^®^ qPCR assays varied between 93 % and 106.7%. The analytical sensitivity or limit of detection of the assays varied between 0.6 and 1.4 copies/µl (Table 4). The specificity in G1_3_qPCR, G4_qPCR and G5_qPCR was 100 %. G6_10_qPCR cross-reacted with *E. ortleppi* (G5) samples (Table 3) and required a duplex assay design with G5_qPCR for diagnostic purposes.

**Table 4:**
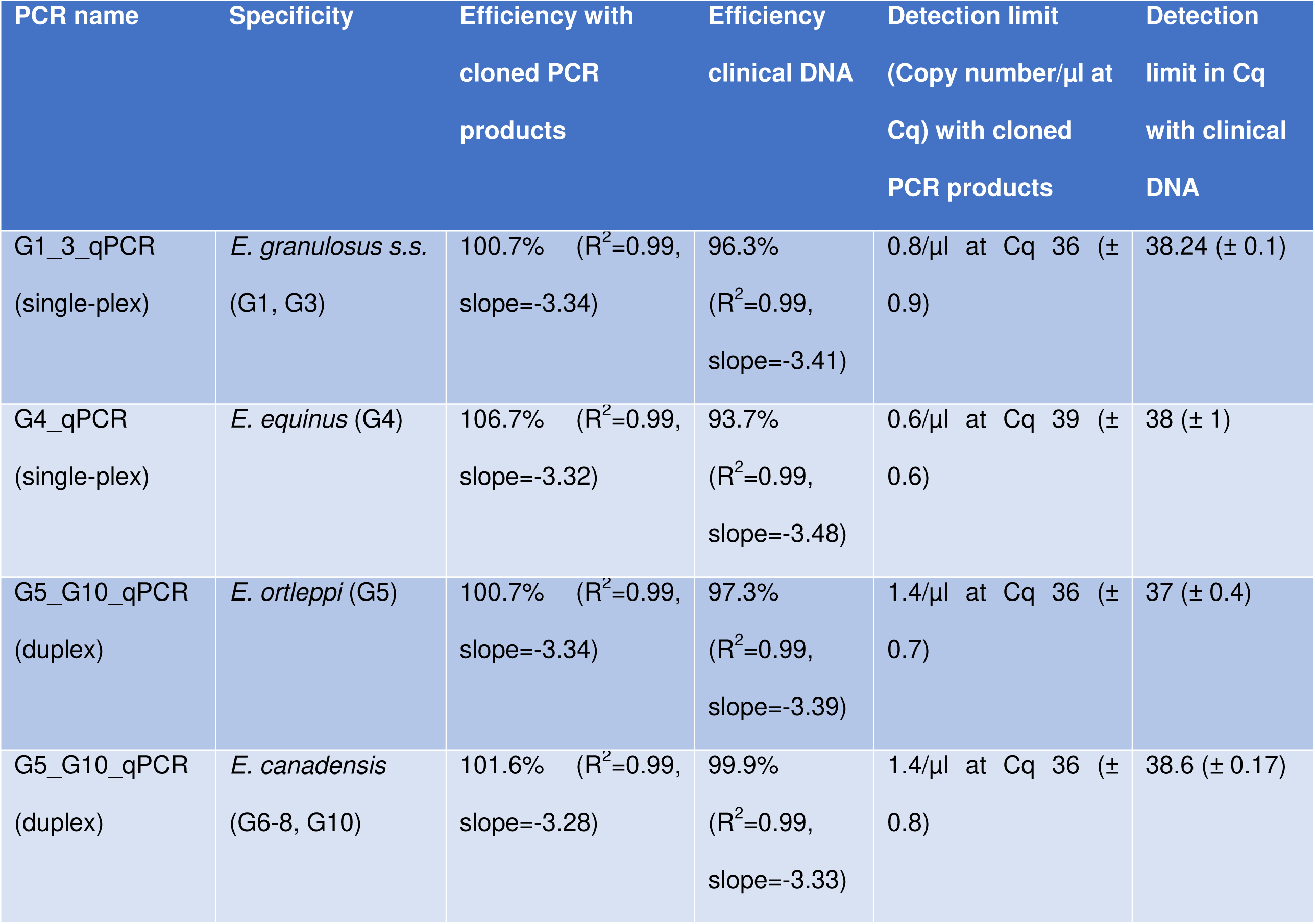
Analytical sensitivity and efficiency of developed TaqMan^®^ qPCRs.

**Figure 5:**
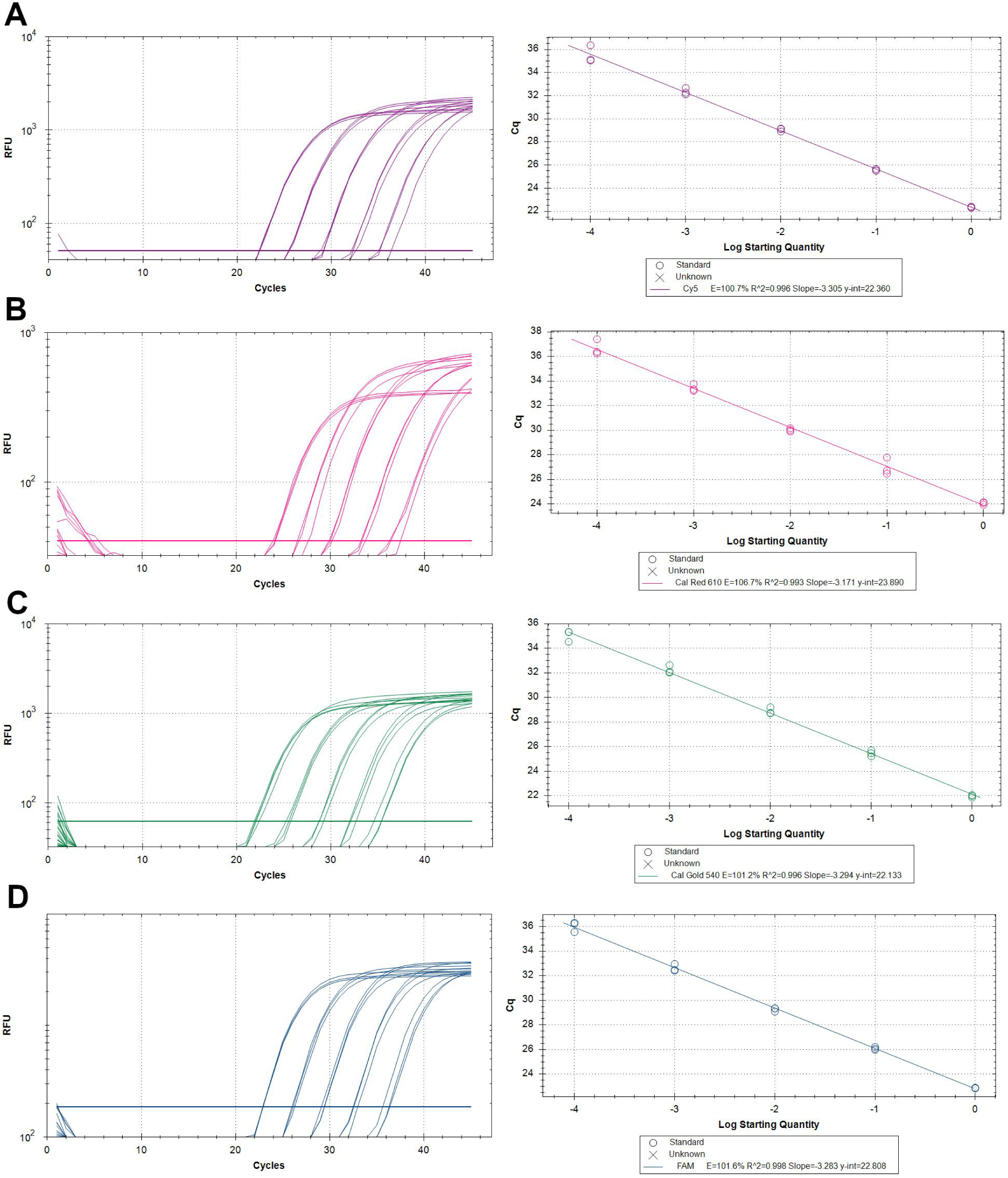
Fluorescence curves of serially diluted specific plasmid DNA samples analysed by TaqMan® qPCR. **A**: G1_3_qPCR: *E. granulosus s*.*s*., **B:** G4_qPCR: *E. equinus*, **C**: G5_10_qPCR: *E. ortleppi*, and **D:** G5_10_qPCR: *E. canadensis*. RFU: relative fluorescence units. Cq: quantification cycle.

In summary, sequence-specific DNA probe-based TaqMan^®^ qPCR assays were established that identified four species within *E. granulosus s*.*l*. in reference samples. These samples were further differentiated from other *Echinococcus* and *Taenia* species. (Table 3).

### Detection of *E. granulosus s*.*l*. species in faecal samples

The reference DNA samples used in this study for the development of qPCR assays that differentiate four *Echinococcus* species was derived from cyst wall material of metacestodes isolated from infected hosts. While cysts may serve as a sample matrix for the detection of *E. granulosus s*.*l*. species, it is diagnostically relevant that the assays also amplify target DNA extracted from other relevant matrices such as faecal matter. To test the performance of the TaqMan^®^ qPCR assays when used with faecal samples, fox faeces were spiked with serially diluted reference DNA to simulate the testing of faeces from definitive hosts infected with members of *E. granulosus s*.*l*.. A known quantity of standardised, heterologous plasmid DNA with matching primers and probe was included as an internal control [IC] in the qPCR mixture (31) to control for PCR inhibition by faecal factors. Inclusion of the IC also allowed testing the assays in duplex (G1_3_qPCR or G4_qPCR in combination with IC-qPCR) and triplex (G5_G10_qPCR and IC-qPCR) TaqMan^®^ qPCR formats (see Table 1). All four assays targeting *E. granulosus s*.*s*. (G1-G3), *E. equinus* (G4), *E. ortleppi* (G5) and *E. canadensis* (G6-8, G10) respectively, amplified the specific tapeworm DNA region from each analysed faecal sample type (Table 5). Combining *E. ortleppi, E. canadensis* as well as IC primers and respective probes in a triplex qPCR differentiated *E. ortleppi* (G5) and *E. canadensis* (G6-8, G10) DNA in faecal samples (Table 5). Probe recognition of IC DNA and quantification showed that no inhibition of the amplification process had occurred in these faecal samples and that the selected primer probe mixtures performed reliably with samples prepared in a matrix of faecal matter under duplex or triplex qPCR conditions. The Cq values generated from the re-extracted tapeworm DNA from faeces were much higher than those from the non-faecal control samples i.e. DNA extracted from hydatid cyst material. This suggests that the quantities of re-extracted tapeworm DNA from faeces were much lower than from non-faecal control samples, possibly due to loss of DNA during the faecal DNA extraction process (Table 5).

**Table 5:** Detection of *E. granulosus s*.*l*. species in faecal samples by duplex and triplex TaqMan^®^ qPCRs

| PCR Name | Specificity | 1:100§ | 1:100* | 1:1000* | 1:10000* | 1:100000* |
| --- | --- | --- | --- | --- | --- | --- |
| G1_3_qPCR+IC_qPCR<br>(duplex) | G1, G3 | 24.5 | 30.1 (±0.6) | 35.2 (±1) | neg. | neg. |
| G4_qPCR+IC_qPCR<br>(duplex) | G4 | 20.2 | 27.4 (±0,5) | 30.7 (±1) | 35.8 (±0.06) | neg. |
| G5_10_qPCR+IC_qPCR<br>(triplex) | G5 | 21,37 | 30.3 (±0.17) | 32.4 (±0.22) | neg. | neg. |
| G5_10_qPCR+IC_qPCR<br>(triplex) | G6-10 | 24.2 | 31.2 (±0.02) | 32.6 (±0.22) | neg. | neg. |

§Cq values from 1:100 pre-diluted DNA, extracted from cyst material
\*Cq-Values from re-extracted DNA prepared by logarithmic dilution of the corresponding DNA, used to spike faecal samples (± standard deviation)

Taken together, we developed four sequence specific-DNA probe-based qPCR assays that allow differentiation of *E. granulosus s*.*l*. species detected in DNA samples derived from cyst material or spiked faecal matter, namely *E. granulosus s*.*s*. (G1-G3), *E. equinus* (G4), *E. ortleppi* (G5) and *E. canadensis* (G6-8, G10).

## Discussion

Cystic echinococcosis is a globally important parasitic zoonosis that requires well-informed prevention and control measures based on reliable and efficient diagnosis of the various causative agents (32, 33).

By designing TaqMan^®^-qPCR-probes that directly identify polymorphic genome regions among the four most important species of the *E. granulosus s*.*l*. complex, we simplify and enhance current diagnostic procedures in multiple ways. The advantages include increased sensitivity, increased specificity, the ability to quantify sample DNA, reduced time-cost to achieve a diagnostic result and potentially reduced processing and equipment costs due to focussing on a single technology. Taken together, these methodological properties should significantly facilitate the process of establishing diagnostic capacities for the detection of *E. granulosus s*.*l*. in field laboratory settings and also facilitate higher sample throughput for epidemiological studies (25, 26).

We discriminate the individual CE agents diagnostically by targeting genetic variability of the mitochondrial genome. Using a bioinformatics approach, we first identified polymorphic regions in the genes *Cox1, Cox3* and *Nad5* (Figure 1), which have previously emerged as suitable targets for *E. granulosus s*.*l*. genotyping (11, 34). Selected primer-probe combinations for all but one species were specific, as the single-plex qPCR system for *E. canadensis* (G6-8, G10) cross-reacted with *E. ortleppi* (G5) samples (Table 3). One obvious possible explanation is the close relationship of these species, resulting in a low degree of polymorphism between *E. ortleppi* and *E. canadensis* in the *Cox3* region targeted by G6-10 primers and probe. However, the non-specific G6-10-TaqMan^®^-probe directed at *Cox3* differed from its corresponding G5 sequence by four nucleotides (G>A, G>A, C>T, C>T), whereas the specific G5-*Nad5*-probe differed only by three nucleotides when compared to its corresponding G6-10-sequence (C>T, C>T, G>A) (Figure 1). Therefore, it appears that nucleotide polymorphism alone is not sufficient to determine probe specificity in this context and that other single stranded DNA-affinity mechanisms, such as steric nucleotide relations (35), interact to overcome the mismatching probe-sequence in the G6-10 qPCR system. Our cross-reactivity observations are also consistent with genome phylogeny studies demonstrating a close relationship between *E. ortleppi* and *E. canadensis* (4, 13). Born from necessity, we therefore developed a duplex qPCR format, which now enabled successful G5 and G6-10 diagnosis for both samples in a single, parallel step, thus highlighting the value of multiplex qPCR diagnosis for *Echinococcus* species in the future.

Whilst phylogenetic relationship may contribute to diagnostic cross-reactivity, testing of *E. granulosus s*.*s*. haplotypes, which do not precisely fit with sequences of G1 and G3, but clearly belong to that species (here named Gx), showed that our assay correctly classified a “non-conventional” echinococcal isolate (Figure 2). Phylogenetic studies of *E. granulosus s*.*s*. isolates found that a large proportion of the identified haplotypes were not homologous with G1 and G3 in the classical sense of the G-nomenclature, yet they clearly belonged to the *E. granulosus s*.*s*. cluster (4, 13). Here, we classify the G_x_-isolate that could not be assigned to the conventional G1-3 genotypes previously (4, 13) as *E. granulosus s*.*s*., which indicates a broad applicability of the G1_3_qPCR assay in the context of the natural genetic diversity in this subgroup (for an alternative interpretation of genotypes G1 and G3 see (10)).

The described qPCR system includes all species and genotypes of *E. granulosus s*.*l*. except *E. felidis*. However, this species, which is not known to be zoonotic, has a very fragmented distribution range in sub-Saharan Africa as it apparently depends on the presence of its principle definitive host, the lion. In any case, the genetic structure of African echinococcosis seems to be more complex than that found elsewhere and is in need of further research. This is illustrated by the presence of a highly divergent zoonotic genotype, G-Omo, in north-eastern Africa. This genotype was only tentatively retained in *E. granulosus s*.*s*. pending further epidemiological information (6). Interestingly, DNA of G-Omo did not react in the *E. granulosus s*.*s*. qPCR (as *E. felidis*, which belongs to the same species cluster), which supports a separate identity of this taxon. Likewise, *E. canadensis* may in future have to be split in two species, namely the wildlife transmitted G8 and G10, and the largely domestically transmitted G6/7 genotype group(36). This will not diminish the value of the qPCR, as both genotype cluster are allopatrically distributed and occur in very different epidemiological settings, with only a limited area of possible overlap in the Asian part of the Russian Federation (6).

The TaqMan^®^-probe-based qPCR assays developed in this study show good efficiency, analytical specificity as well as methodical and diagnostic sensitivity when used with DNA obtained from cysts. Therefore, they are suitable for diagnosis of *Echinococcus* species in intermediate hosts, including humans. Our findings in spiked faecal samples suggest that the duplex qPCR should also be applicable for the detection of *E. granulosus s*.*l*.-egg DNA in definitive hosts. Detection and genotyping of *Echinococcus* spp. DNA in faecal samples would be particularly useful when examining larger populations of living animals in the field for epidemiological studies,. Whilst we characterised the analytical sensitivity of the assays by using cloned amplicons (Table 4, Figure 4 and Figure 5), their methodical and diagnostic sensitivity with faecal samples containing parasite eggs remains to be determined. A similar standardised, cloned-amplicon-based approach could be used to quantify target DNA in sample matrix of interest when using our assays. For instance, we observed an approx. 100-fold loss of specific DNA qPCR detection in spiked faecal samples (approx. 7 Cq-values) when compared to similar DNA quantities analysed in cyst material (Table 5). Inclusion of the IC-plasmid internal control suggested that the DNA loss occurred at the extraction step, as the amplification in faecal and non-faecal samples was comparable (Table 4). This observation further showed that faecal inhibitors known to potentially interfere with PCR amplification (28), had no noticeable effect in our qPCR system. However, faecal samples from other animals could harbour another combination of the inhibitory components, which may negatively influence the amplification process of the assays (28).

We anticipate that single-step genotyping techniques for *Echinococcus granulosus s*.*l*. complex diagnosis by DNA-probe-based qPCR will complement the available methods, improve case reporting on the genotype-subgroup-level, advance our knowledge on the epidemiology of these parasites and ultimately support effective control of CE.

## Acknowledgements

We would like to thank Alrik-Markis Kunisch and Susanne Zahnow for technical assistance. We also thank Dr. Bernd Hoffmann for discussions on the study design and Dr. Johanna Dups-Bergmann for critical reading of this manuscript. Furthermore we would like to thank Hasmik Gevorgian for providing *E. granulosus s*.*s*. cyst material from Armenia, Yitagele Terefe for providing *E. granulosus s*.*s*. (Gx), *T. saginata* and *T. hydatigena* cyst material from Ethiopia, Ortwin Aschenborn for providing *E. equinus* (G4) cyst material from Namibia, Cecilia Mbae for providing *E. ortleppi* (G5) and *E. canadensis* (G6) from Kenya, Fredrick Banda for providing of *E. ortleppi* (G5) cyst material from Zambia, Sergey Konyaev for providing of *E. canadensis* (G8) cyst material from Russia, Liudmila Kokolova for providing of *E. canadensis* (G10) from Russia and Daniel Woldeyes for providing of *E. canadensis* (G8) cyst material from *E. cf. granulosus (G*_*omo*_) from Ethiopia. This work was supported by funding from the European Union’s Horizon 2020 Research and Innovation programme under grant agreement number 773830: One Health European Joint Programme (MEME project; https://onehealthejp.eu/jrp-meme/).

## Declaration

The study described is original and is not under consideration by any other journal. All authors approved the final manuscript and its submission. The authors declare that they have no conflict of interest.

